# High frequency visual stimulation primes gamma oscillations for visually-evoked phase reset and enhances spatial acuity

**DOI:** 10.1101/2020.09.29.319467

**Authors:** Crystal L. Lantz, Elizabeth M. Quinlan

## Abstract

The temporal frequency of sensory stimulation is a decisive factor in the bidirectional plasticity of perceptual detection thresholds. However, surprisingly little is known about how distinct temporal parameters of sensory input differentially impact neuronal, circuit, and perceptual function. Here we demonstrate that brief repetitive visual stimulation is sufficient to induce long-term plasticity of visual responses, with the temporal frequency of the visual stimulus determining the location and generalization of visual response plasticity. Brief repetitive low frequency stimulation (LFS, 2 Hz) is sufficient to induce a visual response potentiation that is exclusively expressed in layer 4 in response to the familiar stimulus. In contrast, brief, repetitive high frequency stimulation (HFS, 20 Hz) suppresses the activity of fast-spiking interneurons and primes ongoing gamma oscillatory rhythms for visually-evoked phase reset. Accordingly, visual stimulation subsequent to HFS induces non-stimulus specific visual response plasticity that is expressed in all cortical layers. The generalized visual response enhancement induced by HFS is paralleled by an increase in visual acuity measured by improved performance in a visual detection task.

## Introduction

Enduring bidirectional changes in synaptic strength induced by specific frequencies of synaptic stimulation are a primary mechanism for information storage in neural circuits. A wealth of data from brain slices demonstrates that high frequency stimulation rapidly induces long-term potentiation (LTP) of excitatory synapses, while low frequency stimulation induces synaptic long-term depression (LTD; Bliss and Lømo, 1973; Dudek and Bear, 1992; Kirkwood et al., 1996; Mayford et al., 1995). Repetitive stimulation *in vivo* in animal models reveals a similar dependence of bidirectional synaptic plasticity on stimulation temporal frequency, with synaptic depression and potentiation induced by low and high frequency stimulation respectively (Nabavi et al., 2014; Rodríguez-Durán et al., 2017; O’Riordan et al., 2018). Changes in synaptic strength evoked by specific temporal patterns of stimulation are also the foundation for frequency-dependent changes in neuronal activity and perception induced by transcranial stimulation. Experiments in human subjects utilizing sensory stimulation with temporal frequencies associated with LTP induction demonstrate rapid enhancement of response amplitudes and lowering of perceptual discrimination thresholds in visual, auditory and somatosensory systems (Beste et al., 2011; Brickwedde et al., 2020; Marzoll et al., 2018; Pegado et al., 2016; Ross et al., 2008; Sanders et al., 2018). This enhancement of sensory responses in humans and animal models induced by non-invasive sensory stimulation has been shown to require NMDA receptor activation, mimicking the induction requirements for LTP (Clapp et al., 2006; Dinse et al., 2003).

Sensory response amplitudes and perceptual detection thresholds reflect the interaction between stimulus-evoked neuronal responses and ongoing fluctuations in the cortical local field potential (LFP; Sauseng et al., 2007; Xing et al., 2012). In the mouse primary visual cortex (V1), the power of multiple layer 4 oscillatory bands increases in response to a familiar stimulus, suggesting that visual stimulus familiarity can be encoded by changes in oscillatory power (Kissinger et al., 2018). In concert, the magnitude and perception of incoming visual stimuli are regulated by the phase of cortical oscillations (Kim et al., 2007). The temporal frequency of incoming auditory and somatosensory stimulation can also impact sensory perception through entrainment of ongoing rhythmic activity (Brickwedde et al., 2020; ten Oever et al., 2017). Indeed, increased power and synchronization of high frequency cortical oscillations underlies improvements in stimulus-detection and memory encoding by attention and motivation (Jutras et al., 2009; Montgomery and Buzsáki, 2007; Lisman, 2010). Thus changes in the time-locked evoked response, as well as the magnitude and phase of ongoing cortical oscillations, are candidate mechanisms for sensory-evoked synaptic plasticity (Brickwedde et al., 2019; Howe et al., 2017; Park et al., 2016).

Long-lasting enhancement of visual responses, that emerge slowly and require NMDA receptor activity, are also induced in mouse V1 following daily repetition of low frequency visual stimulation (LFS; Frenkel et al., 2006). This experience-dependent response plasticity is revealed as an increase in the amplitude of the VEP recorded in layer 4 of V1, and an increase in the peak firing rate of regular-spiking (RS) neurons, in response to a familiar visual stimulus (Cooke et al., 2015). Response potentiation induced by daily LFS is highly selective for the orientation, contrast and spatial frequency of the visual stimulus used for induction. Indeed, response potentiation is not observed following rotation of the orientation of the visual stimulus by as little as 5 degrees (Cooke and Bear, 2010). The time course and selectivity of visual response potentiation shares many similarities with perceptual learning (Poggio et al., 1992; Zaehle et al., 2007). Additionally, the response to daily LFS is occluded by potentiation of thalamo-cortical synapses, and requires sleep for expression, indicating similar induction and consolidation requirement as LTP (Cooke and Bear, 2010; Aton et al., 2014). Suppression of parvalbumin expressing fast spiking (FS IN) output following daily LFS, via optogenetic silencing or the NMDAR antagonist ketamine, decreases the selectivity of visual response potentiation (Kaplan et al., 2016). Moreover, genetic deletion of the immediate early gene neuronal pentraxin 2 (NPTX2; aka NARP), which is highly enriched at excitatory synapses onto parvalbumin-positive interneurons, inhibits the induction of visual response potentiation induced by a single bout of LFS (Chang et al., 2010; Gu et al., 2013; O’Brien et al., 1999; Xu et al., 2003). High temporal frequency visual stimulation (HFS, 20 Hz) rescued visual response potentiation in NPTX2^-/-^ mice (Gu et al., 2013).

Although the temporal frequency of sensory stimulation plays a crucial role in the induction of bidirectional plasticity, little is known about how stimulus temporal frequency recruits distinct neuronal, circuit and perceptual changes. Here we assess the impact of the temporal frequency of repetitive visual stimulation on long-lasting changes in the mouse visual system. We find that a single bout of LFS is sufficient to induce an increase in the magnitude of visual responses that is restricted to layer 4 and the familiar visual stimulus. In contrast a single bout of HFS induces visual responses plasticity throughout V1 in response to familiar and novel stimuli. HFS induces a long-lasting suppression of the output of FS INs and sensitizes cortical gamma oscillations to phase reset by all subsequent visual stimuli. This general enhancement of visual response strength following HFS is paralleled by improved visual acuity revealed by performance in a visual detection task.

## Methods

### Animals

Experiments utilized equal numbers of male and female adult (postnatal day 60-90) C57BL/6J mice (Jackson Lab, Bar Harbor, ME). Subjects were housed on a 12:12 hour dark:light cycle with food and water *ad libitum*, and experiments were initiated ~6 hours into the light phase. All procedures conformed to the guidelines of the University of Maryland Institutional Animal Care and Use Committee. Sample sizes were determined by power analysis of previous studies quantifying the effect of visual experience on visual response amplitudes.

### Electrophysiology

House-made 1.2 mm length 16-channel shank electrodes were implanted into binocular V1 (from Bregma: posterior, 2.8 mm; lateral, 3.0 mm; ventral, 1.2 mm), anesthetized with 2.5% isoflurane in 100% O_2_, as described (Bridi et al., 2018; Murase et al., 2017). Subjects received a single dose of carprofen (5 mg/kg, SQ) for post-surgical analgesia after the return of the righting reflex. One week after surgery and one day before electrophysiological recordings, subjects were habituated for 45 minutes to head restraint. Broadband electrophysiological data was collected from awake head-fixed mice, using RZ5 bioamp processor and RA16PA preamp (Tucker Davis Technologies, TDT), and multiunit waveforms were sorted into single units using an automatic Bayesian clustering algorithm in OpenSorter (TDT) as described (Murase et al., 2017). Single units (SUs) were processed in MATLAB and classified as regular spiking neurons(RS, presumptive excitatory) or fast spiking interneurons(FS IN, presumptive inhibitory) based on waveform slope 0.5 msec after the trough, time between trough and peak, and the ratio of trough to peak height (Niell and Stryker, 2008). VEPs and SUs were assigned to cortical layer based on LFP waveform shape, and current source density calculated with single site spacing from the laminar array (Guo et al., 2017; Mitzdorf, 1985).

### Visual Stimulation

Visual stimuli were presented using MATLAB (Mathworks) with Psychtoolbox extensions (Brainard, 1997; Pelli, 1997). Prior to visual stimulation each day mice passively viewed a grey 26 cd/m^2^ screen for 200 seconds (spontaneous). Visually evoked responses were recorded in response to 200 x 1s trials of 0.05 cycles per degrees, 100% contrast, square-wave gratings reversing at either 2 Hz (LFS) or 20 Hz (HFS) at various orientations. VEPs and SUs were recorded 24 hours after initial LFS or HFS visual stimulation, in response to familiar and novel stimulus orientations and spatial frequencies (200 x 1s trials of 0.05 cycles per degrees, 100% contrast, square-wave gratings reversing at 2 Hz).

### Data Analysis

Spike rates of sorted SUs were calculated as the average firing rate during each 1 second epoch (200 trials). Peristimulus-time histograms (PSTH) were calculated for each SU using 5 ms bins and smoothed with a gaussian kernel (Kissinger et al., 2018). To examine changes in oscillatory power by frequency, PSTHs were z-scored, and filtered from 1 to 100 Hz using a sliding frequency window via a bandpass elliptic filter with a span of 3 Hz in MATLAB. The analytic signal of band-passed PSTHs was calculated using a Hilbert transform, and the absolute value was used to calculate power within each frequency band.

Visually evoked potentials (VEPs) were calculated as the trough to peak amplitude of the average of 1 second LFP epochs during visual stimulation in MATLAB, as described (Murase et al., 2017). To examine changes in oscillatory power by frequency and the reliability of incoming visual stimulation to reset the phase of ongoing oscillations (Inter-Trial Phase Consistency, ITPC, a time locked measure of oscillatory phase), spontaneous and evoked LFPs were z-scored then convolved with complex Morlet wavelets from 1 to 100 Hz using a 3Hz window (Fiebelkorn et al., 2018). The wavelet cycle width varied with filtered frequency (1-10 Hz = 2 cycles, 11-14 Hz = 3 cycles, 15-20 Hz = 4 cycles, 21-100 Hz = 5 cycles). The absolute value of the complex vector was used to calculate oscillatory power. Power was averaged between trials for each subject, and activity is reported as percent change in power relative to spontaneous activity recorded on the first day prior to experimental manipulation (experimental – baseline / baseline x 100). Averaged percent change in evoked power was calculated during the 100-200 ms post stimulus onset, binned oscillatory activity was averaged from this window. Oscillatory bins were defined as: delta 1-4 Hz, theta 4-8 Hz, alpha 8-13 Hz, beta 13-30 Hz, and gamma 30-100 Hz. The second half of the complex vector was normalized, averaged and the absolute value was used to calculate ITPC. ITPC was binned by oscillatory frequency. Utilizing the calculated oscillatory phase of the LFP, the phase of each frequency for each single unit was calculated then averaged as Spike-Phase Consistency.

### Behavior

Psychophysical measurements of spatial acuity were assessed with performance in a 2 alternative forced choice visual detection task. Task training and testing utilized a Bussey-Saksida Touch Screen Chamber (Lafayette Instruments; Horner et al., 2013), with custom plexiglass inserts. An opaque insert divided the LCD touch screen into two vertical halves, for simultaneous display of the correct and incorrect visual stimuli. A transparent insert, parallel to and 6cm from the touch screen, with 2 5×7 cm swing-through doors, defined the choice point for the calculation of visual stimulus spatial frequency (Fig 7A). Naïve adult mice were trained to associate a high contrast (100%), low spatial frequency (0.05 cycles/degree), 45° sinusoidal grating (positive stimulus) with positive reinforcement (strawberry milk and tone, 3kHz, 0.5s) and a grey image of equal luminance (32 cd/m2; negative stimulus) with negative reinforcement (tone, 400 Hz, 2s). Subjects were food deprived for 20 hours a day, with 4 hours of *ad libitum* food access at the end of each training session. A training session consisted of 30 trials, or 45 minutes. To begin a trial, the subject nose poked in the illuminated liquid reward tray at the rear of the chamber, to trigger the presentation of positive and negative visual stimuli. Contact with the touch screen turned the visual stimulus off. Choosing the positive stimulus resulted in a liquid reward, choosing the negative stimulus resulted in negative reinforcement and a 30 second timeout period. Trials were separated by a 10 second intertrial interval, followed by illumination of the liquid reward tray to signal the beginning of a new trial. Criterion was defined as 25/30 correct trials within 45 minutes = 83% correct). After reaching criterion, acuity testing was initiated. In acuity testing, the positive stimulus was rotated to a novel orientation (45°+15°). Following successful completion (≥ 60%) of a block of 10 trials, the spatial frequency was increased incrementally (0.05 cpd steps). The highest spatial frequency with performance of ≥ 70% correct choices is defined as spatial acuity. Following assessment of initial acuity subjects were randomly divided into 2 groups, 50% viewed 200 seconds of LFS, 50% viewed 200 seconds of HFS, at a novel orientation, and were returned to their home cage. 24 hours after visual stimulation, acuity was tested at a familiar stimulus orientation (used for LFS or HFS) and at a novel orientation, with test order randomized. Subjects were then returned to food and water *ad libitum* in the mouse colony.

### Statistics

Statistical analysis was completed using JASP (JASP Stats). Repeated measures ANOVA (RANOVA) was used to compare LFP data arising from 3 time points within the same subject, including VEP, oscillatory power and ITPC, followed by a Bonferroni *post-hoc* when appropriate. We did not assume that signal from the same electrode was the same single unit over multiple days, therefore, unpaired Student’s t-test was used to compare 2 groups and a one-way ANOVA was employed to compare 3 groups, followed by a Tukey *post-hoc* when appropriate. A multivariate ANOVA (MANOVA) with a Bonferroni *post-hoc*, when appropriate, was used to compare oscillatory data consisting 2 time points with multiple frequency bands. To compare the change in oscillatory power within subjects we employed a one-sided Student’s t-test with a Bonferroni correction for multiple comparisons. In text, n is reported as the total number of subjects, followed by the total number of recorded single units. Exact p values are reported in the text, except when p < 0.001.

## Results

### Visual stimulus frequency controls plasticity of visual responses

To examine the impact of the temporal frequency of repetitive visual stimulation on visual response plasticity, we examined the magnitude of visually-evoked potentials (VEPs) throughout the depth of the primary visual cortex (V1). Head-fixed awake mice viewed 200 x 1s trials of square-wave gratings (0.05 cycles per degrees, 100% contrast, 60° orientation) delivered at low frequency (2 Hz) or high frequency (20 Hz, Fig. 1A&C). Long lasting visual response potentiation was induced by both protocols, while the location and generalization of visual response potentiation was determined by the visual stimulus temporal frequency. 24 hours after a single bout of low frequency stimulation (LFS), the amplitude of the layer 4 VEP was significantly increased in response to familiar (60°), but not novel (150°) visual stimulus orientations, mimicking the specificity of stimulus selective response potentiation induced by LFS over multiple days (Frenkel et al., 2006; n = 16, RANOVA_(df, 2, 15)_, Bonferroni *post-hoc*, F = 9.13, p < 0.001; initial v. familiar: p = 0.023, Fig. 1B).

**Figure 1.**
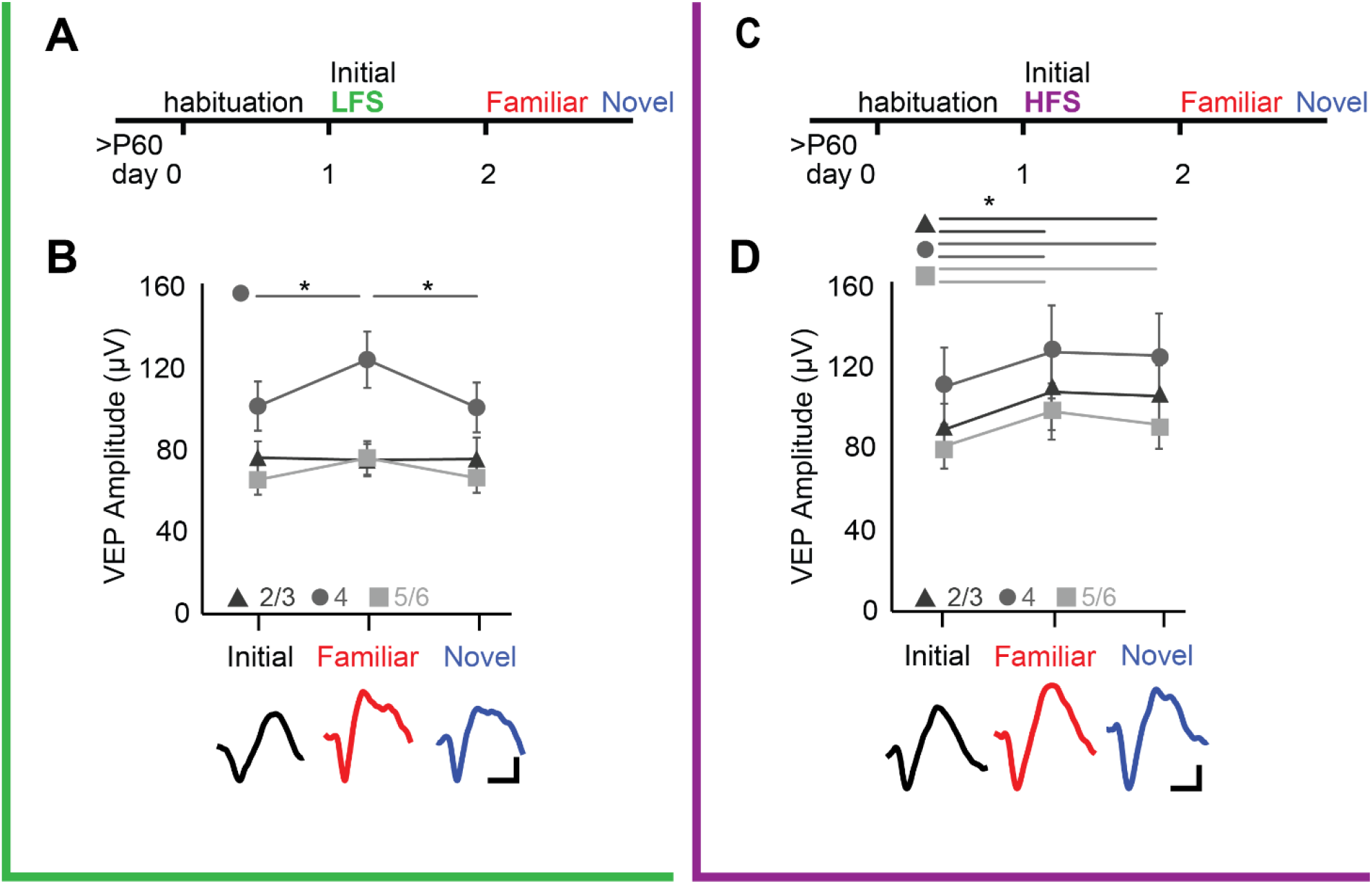
LFS and HFS differentially impact the location and generalization of visual response potentiation. A) Experimental timeline: naïve subjects received low frequency visual stimulation (LFS, green: 200 presentations of 0.05 cpd, 100% contrast gratings, 60° orientation, 2 Hz). The initial response is reported as baseline VEP. After 24 hours, VEPs were evoked by familiar and novel visual stimulus orientations presented at 2 Hz. B) Significant increase in VEP amplitude in layer 4 in response to familiar, but not novel, stimulus orientations (RANOVA _(df, 2, 15)_, F = 9.13, p < 0.001; * = Bonferroni *post hoc* p < 0.05; n=16 subjects). C) Timeline, as in A, except that naïve subjects received high frequency visual stimulation (HFS, purple: 200 presentations of 0.05 cycle per degree, 100% contrast grating, 30° orientation, 20 Hz) after initial assessment of baseline VEP. After 24 hours, VEPs were evoked by familiar and novel visual stimulus orientations presented at 2 Hz. D) Significant increase in VEP amplitudes in layers 2/3, 4, and 5/6 in response to familiar and novel stimulus orientations (RANOVA _(df, 2, 10)_, layer 2/3: F = 5.25, p = 0.015, layer 4: F = 7.31, p = 0.004; layer 5/6: F = 11.41, p < 0.001; * = Bonferroni *post hoc* p<0.05; n = 11 subjects).

In contrast, 24 hours after a single bout of high frequency stimulation (HFS), the amplitude of VEPs throughout all layers of V1 were significantly increased in response to the familiar (60°) and novel (150°) visual stimulus orientations (n=11, RANOVA _(df, 2, 10)_, Bonferroni *post-hoc*, layer 2/3: F = 5.25, p = 0.015, initial v. familiar p = 0.023, initial v novel p = 0.032; layer 4: F = 7.31, p = 0.004, initial v. familiar p = 0.038, initial v. novel: p = 0.005; layer 5/6: F = 11.41, p < 0.001, initial v. familiar: p = 0.005, initial v. novel: p = 0.022, Fig. 1D). A similar potentiation of VEP amplitudes that generalized to novel stimuli was induced if HFS stimulation preceded or followed assessment of baseline VEP amplitudes (Between subjects RANOVA _(df 1,12)_ layer 2/3: F = 0.001, p = 0.97, layer 4: F = 0.010, p = 0.92, layer 5: F = 0.00003, p = 0.995, data not shown). HFS also induced an increase in VEP amplitudes evoked by visual stimuli of novel spatial frequencies (Fig. S1). Thus, the response plasticity induced by a single bout of LFS is localized and stimulus-specific, and the response plasticity induced by HFS is global and generalizes to novel stimuli.

### Visual stimulus frequency acutely modulates oscillation power, phase, and LFP-Spike coupling in V1

The amplitude of the VEP reflects the interaction between stimulus-evoked neuronal responses and ongoing fluctuations in the cortical local field potential (LFP). Changes in VEP amplitude could therefore reflect changes in the power or phase of LFP oscillations. To ask how the power of LFP oscillations were impacted during LFS and HFS, we normalized the absolute value of the complex Morlet wavelet convolved LFP during visual stimulation to pre-stimulation spontaneous activity (equal luminant grey screen; 26 cd/m^2^). During LFS, low frequency (alpha and beta bands) LFP oscillatory power was significantly increased in all cortical layers (n = 16, one-sided t-test. Layer 2/3: α: t = 4.44, p < 0.001, β: t = 4,39, p < 0.001. Layer 4: α: t = 2.37, p = 0.015, β: t = 2.31, p = 0.024. Layer 5: α: t = 4.07, p < 0.001, β: t = 2.27, p = 0.019, Fig. 2A). Similarly, during HFS, low frequency (delta, alpha and beta bands) power was significantly increased in all layers (n=11, one-sided t-test, layer 2/3; δ: t = 2.42, p = 0.019, α: t = 3.21, p = 0.005, beta, t = 3.57, p = 0.003; layer 4; δ: t = 2.11, p = 0.031, α: t = 2.94, p = 0.008, β: t = 2.82, p = 0.009; layer 5; δ: t = 2.31, p = 0.023, α: t = 1.93, p = 0.042, β: t = 1.92, p = 0.043; Fig. 2C).

**Figure 2.**
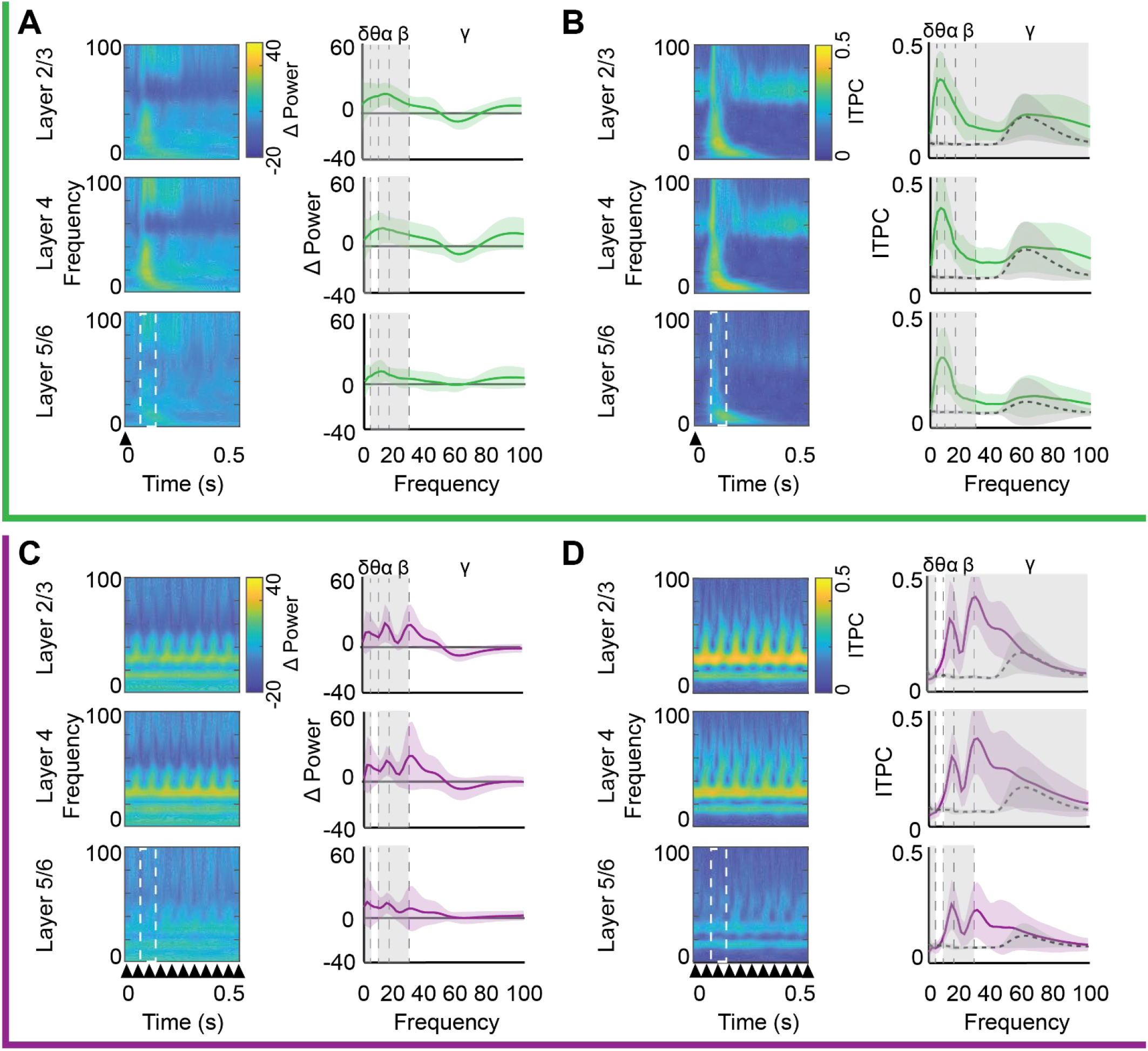
Distinct acute impact of LFS and HFS on oscillatory power and evoked phase reset. Top, green, during LFS: A) Left; average oscillatory power (heat map) from 0-100 Hz (3 Hz bins; y axis) over time (x axis) by cortical layer during LFS. Power was normalized to spontaneous activity in response to 26 cd/m^2^ grey screen. Right; significant increase in average α and β power (y axis) across all cortical layers, in δ power in layers 2/3 and 4, and in θ power in layers 2/3 and 5/6 during LFS (one-sided t-test, Grey highlight = p < 0.05, n = 16). Power binned by frequency band (δ: 1-4, θ: 4-8, α: 8-13, β: 13-30, γ: 30-100Hz). B) Left; average intertrial phase consistency during LFS (ITPC; heat map) from 0-100 Hz (in 3 Hz bins; y axis) over time (x axis), trial averaged. Right; a significant increase in ITPC at all frequencies in layer 2/3 (MANOVA_(df, 1, 5)_, F = 15.94, p < 0.001), and an increase in all frequencies below gamma in layers 4 (MANOVA_(df, 1, 15)_, F = 15.98, p < 0.001) and 5/6 (MANOVA_(df, 1, 15)_, F = 11.10, p < 0.001) during LFS (solid line), relative to spontaneous activity (dashed line). Grey highlight = Bonferroni *post hoc* p < 0.05; n = 16 subjects. Bottom, purple, during HFS: C) Left; average oscillatory power (heat map) from 0-100 Hz (3 Hz bins; y axis) over time (x axis) by cortical layer during HFS, normalized as in A. Right; HFS evoked a significant increase in average δ, α and β power in all cortical layers and in θ power in layer 2/3 (one-sided t-test, Grey highlight = p < 0.05, n = 11). D) Left; average inter-trial phase consistency during HFS (ITPC; heat map) from 0-100 Hz (in 3 Hz bins; y axis) over time (x axis), trial averaged as in B. Right; HFS (purple line) induced significant changes in ITPC in all cortical layers, including a significant increase in visually driven phase reset in α, and β oscillations (layer 2/3: MANOVA_(df, 1, 10)_, F = 14.14, p < 0.001, layer 4: MANOVA_(df, 1, 10)_, F = 12.284, p < 0.001, layer 5: MANOVA_(df, 1, 10)_, F = 9.95, p = 0.0017), and a significant decrease in the δ oscillation in all layers, relative to spontaneous activity (dashed line). Grey highlight = Bonferroni *post hoc* p < 0.05; n = 11 subjects.

To ask how the temporal frequency of visual stimulation impacted the phase of ongoing LFP oscillations, we convolved the LFP signal with a complex Morlet wavelet and calculated the angle of the resultant complex output. Inter-trial phase consistency (ITPC), which ranges from 0 (if phase was random, and not reset by incoming visual input) and 1 (if phase was reset similarly in all trials), was calculated from the time-locked phase and compared to the ITCP during pre-stimulation spontaneous activity. LFS and HFS had differential effects on phase reset of ongoing LFP oscillations. LFS increased phase reset in low frequencies (delta, theta, alpha, and beta) in all cortical layers and increased gamma phase reset in layer 2/3 (n = 16 subjects, Layer 2/3: MANOVA _(df, 1, 5)_ F = 15.94, p < 0.001, δ: F = 36.41, p < 0.001, θ: F = 75.21, p < 0.001, α: F = 88.20, p < 0.001, β: F = 42.22, p < 0.001, γ: F = 4.277, p = 0.047. Layer 4: MANOVA _(df, 1, 5)_ F = 15.98, p < 0.001, δ: F = 45.01, p < 0.001, θ: F = 56.15, p < 0.001, α: F = 65.05, p < 0.001, β: F = 39.86, p < 0.001. Layer 5/6: MANOVA _(df, 1, 5)_ F = 11.10, p < 0.001, δ: F = 20.59, p < 0.001, θ: F = 51.07, p < 0.001, α: F = 46.05, p < 0.001, β: F = 35.09, p < 0.001; Fig. 2B). In contrast, HFS significantly increased phase reset of intermediate frequencies (alpha and beta) in all cortical layers, and increased gamma reset in layers 2/3 and 4. Interestingly, HFS decreased delta reset in all cortical layers, with no changes observed in the phase of theta oscillations (n = 11 subjects, Layer 2/3: MANOVA _(df, 1, 5)_ F = 10.46, p < 0.001, δ: F = 7.76, p = 0.011, α: F = 16.68, p < 0.001, β: F = 43.33, p < 0.001, γ: F = 10.63, p = 0.004. Layer 4: MANOVA _(df, 1, 5)_ F = 11.26, p < 0.001, δ: F = 10.09, p = 0.005, α: F = 28.28, p < 0.001, β: F = 48.15, p < 0.001, γ: F = 5.60, p = 0.028. Layer 5/6: MANOVA _(df, 1, 5)_ F = 6.46, p = 0.002, δ: F = 5.84, p = 0.025, α: F = 24.25, p < 0.001, β: F = 34.24, p < 0.001; Fig. 2D).

The output of fast spiking cortical interneurons (FS INs) expressing parvalbumin play a fundamental role in the control of sensory evoked LFP amplitudes by influencing the generation of theta rhythms and the power and synchrony of gamma rhythmicity (Cardin et al., 2009; Sohal et al., 2009; Stark et al., 2013). To quantify the pattern and strength of the spiking output of individual FS INs during LFS and HFS we calculated the oscillatory power of the post-stimulus time histogram from single unit activity. Oscillatory power during visual stimulation was normalized to pre-stimulation spontaneous activity. The temporal frequency of the visual stimulus was reflected in the output of FS INs, as LFS increased the power of low (4 - 8 Hz) as well as mid-frequency oscillations (7 - 30 Hz, n = 16 subjects, 23 units. One-sided t-test, θ: t = 2.22, p = 0.016, α: t = 2.04, p = 0.024, β: t = 1.84, p = 0.036; Fig. 3A&B), and HFS increased the power of higher frequencies (13 - 30 Hz, n = 11 subjects, 19 units. One-sided t-test, α: t = 1.81, p = 0.039, β: t = 2.41, p = 0.011. Fig. 3E&F). To quantify the coherence of FS IN activity with on-going LFP oscillations, we utilized the time-locked LFP phase to examine the consistency of FS IN spiking within each oscillatory frequency band during LFS and HFS. LFS significantly increased FS IN firing phase consistency with delta in all cortical layers and theta in layers 2/3 and 4 (Student’s t-test, n = 16 subjects, 22 units, layer 2/3: δ: t = 3.63, p < 0.001, θ: t = 3.66, p < 0.001, layer 4: δ: t = 8.10, p < 0.001, θ: t = 3.87, p < 0.001 layer 5: δ: t = 2.02, p = 0.025, Fig. 3C&D). However, no significant differences in FS IN firing phase consistency were observed during HFS (Fig. 3G&H).

**Figure 3.**
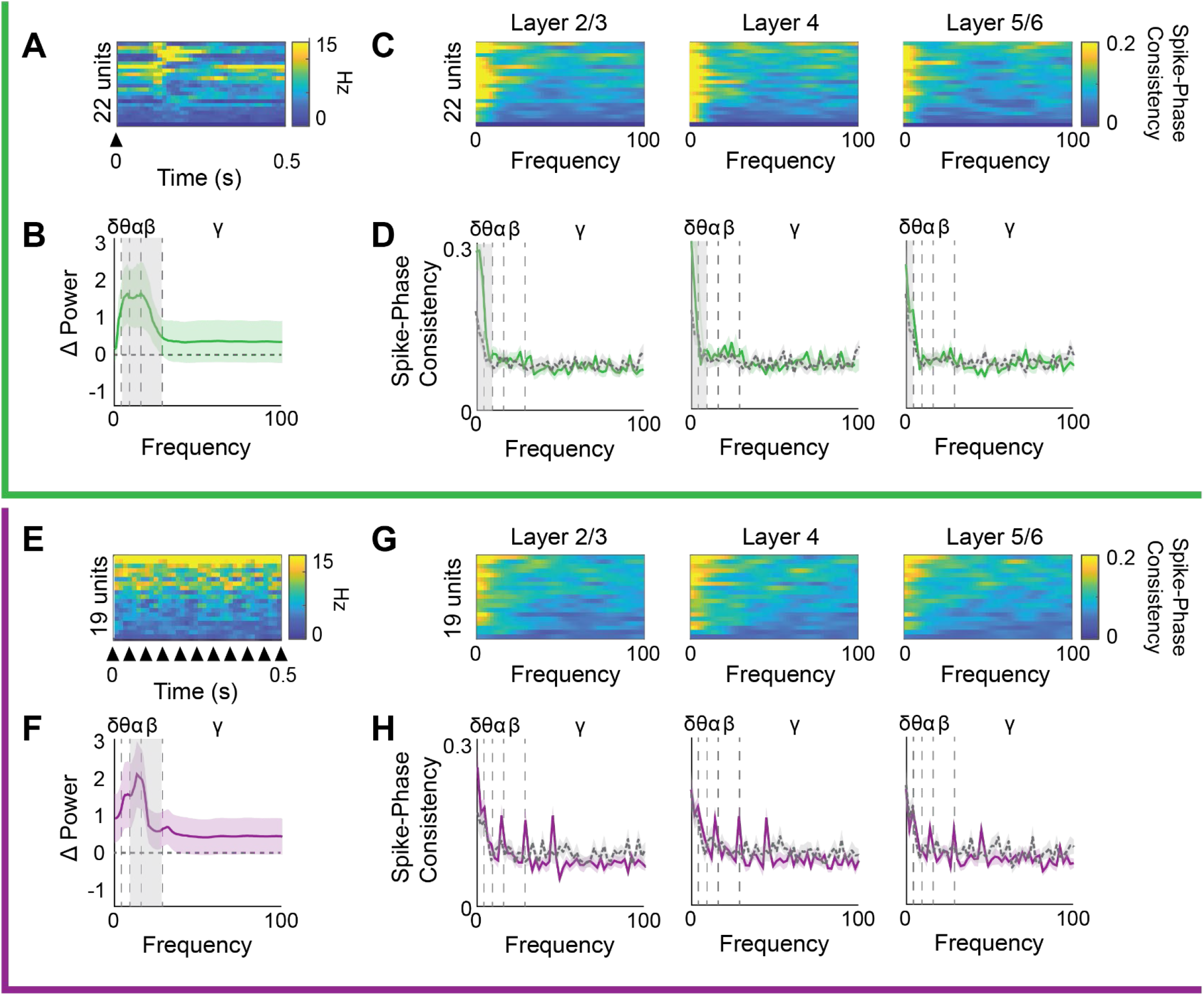
Distinct acute impact of LFS and HFS on FS IN oscillatory power and LFP phase synchrony. Top, green, during LFS: A) Heat map depicting PSTH for each FS IN. B) Average change in oscillatory power of FS INs during LFS, binned by frequency band (δ: 1-4, θ: 4-8, α: 8-13, β: 13-30, γ: 30-100Hz, n = 16 subjects). Significant increase in power of θ, α, and β oscillations in FS IN during LFS (one sided t-test with Bonferroni correction, grey highlight = p<0.05). C) Heat map of FS IN spike-LFP phase consistency during LFS for each FS IN compared to LFP of each layer. D) Average +/- SEM FS IN spike-LFP phase consistency during LFS (green) compared to spontaneous activity (grey), by frequency. LFS increases FS IN spike-phase consistency with δ oscillations in all cortical layers and with θ in layers 2/3 and 4 (Student’s t-test, grey highlight = p < 0.05, n = 16 subjects). Bottom, purple, during HSF: E) Heat map depicting PSTH for each FS IN. F) Average change in oscillatory power of FS IN spiking during HFS, power binned by frequency band (δ 1-4, θ: 4-8, α: 8-13, β: 13-30, γ: 30-100Hz, n = 11 subjects). Significant increase in power of α and β oscillations in FS INs during HFS (one sided t-test, grey highlight = p < 0.05). G) Heat map of FS IN spike-LFP phase consistency during HFS for each FS IN compared to LFP of each layer. H) Average +/- SEM FS INs spike-LFP phase consistency during HFS (purple) compared to spontaneous (grey) by frequency. HFS does not change FS IN spike-phase consistency in any cortical layer (n = 11 subjects).

### Changes in oscillatory power do not predict visual response potentiation

To ask if the visual response plasticity observed in the VEP is reflected in LFP oscillations, we examined the response to the familiar (60° orientation) and novel (150° orientation) visual stimuli 24 hours after LFS or HFS. We utilized the absolute value of the Morlet wavelet convolved LFP in the time window of maximal visually evoked activity (100-200 ms after stimulus reversal). 24 hours after LFS, the familiar visual stimulus induced a significant increase in beta power (13 – 30 Hz) in layers 4 and 5/6 (n = 16, RANOVA _(df, 2, 15)_, Bonferroni *post hoc*. Layer 4: F = 6.51, p = 0.004, initial v. familiar, p = 0.004. Layer 5/6: F = 3.92, p = 0.031, initial v. familiar: p = 0.027; Fig. 4A&B). In contrast, 24 hours after HFS, visual stimuli with familiar and novel orientations triggered a significant decrease in theta power in all cortical layers (n = 11, RANOVA _(df, 2, 10)_ with Bonferroni *post-hoc*. Layer 2/3: F = 8.633, p = 0.002; initial v. familiar, p = 0.015, initial v. novel, p=0.003. Layer 4: F = 8.47, p = 0.002; initial v. familiar, p = 0.016, initial v. novel, p = 0.003. Layer 5/6: F = 7.936, p = 0.003, initial v. familiar, p=0.018; initial v. novel, p = 0.004, Fig. 4 C&D). Additionally, HFS induced a significant decrease in delta power in layers 4 and 5/6 in response to stimuli with the familiar and novel orientations (n = 11, RANOVA _(df, 2, 10)_ with Bonferroni *post-hoc*. Layer 4: F= 10.24, p < 0.001; initial v familiar, p = 0.031, initial v. novel, p < 0.001. Layer 5/6: F = 7.681, p = 0.003, initial v familiar, p = 0.04, initial v novel, p = 0.003, Fig. 4D). Interestingly, the decreases in delta and theta power 24 hours after HFS are also observed in spontaneous activity prior to stimulation with familiar or novel visual stimuli, reflecting the long-term consequence of previous repetitive visual stimulation (Fig. S2D&E).

**Figure 4.**
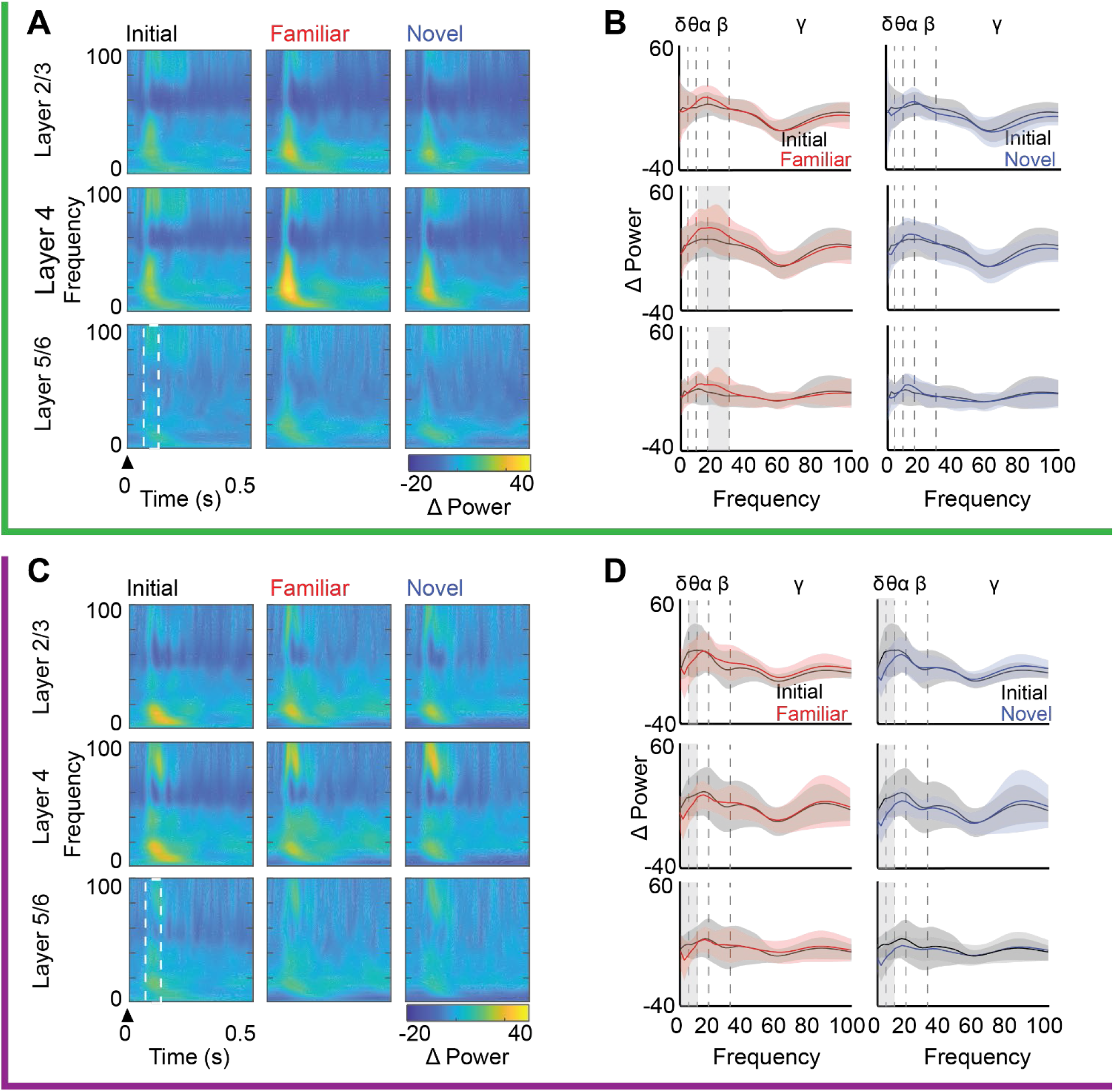
LFS and HFS differentially impact oscillatory power during subsequent visual stimulation. Top, green, after LFS: A) Left; average oscillatory power (heat map) from 0 – 100 Hz (3 Hz bins; y axis) over time (x axis) across cortical layers during initial visual stimulation, and presentation of familiar and novel visual stimuli 24 hours after LFS. Power is normalized to spontaneous activity during viewing of a 26 cd/m^2^ grey screen. Arrowhead indicates stimulus onset, white box indicates time window for assessment of change in oscillatory power (100-200 ms after stimulus onset). B) Average +/- SEM change in oscillatory power, binned by frequency band (δ: 1-4, θ: 4-8, α: 8-13, β: 13-30, γ: 30-100 Hz) during presentation of initial (black), familiar (red), and novel (blue) visual stimuli. In layer 4 and 5/6, a significant increase in average β power in response to familiar (red) but not novel (blue) visual stimuli relative to initial (black, RANOVA _(df, 2, 15)_, layer 4: F = 6.51, p = 0.004; layer 5/6: F = 3.92, p = 0.031). Grey highlight = Bonferroni *post hoc* p < 0.05; n = 16 subjects. Bottom, purple, after HFS: C) Average oscillatory power (heat map) from 0-100 Hz (3Hz bins; y axis) over time (x axis) across cortical layers during initial visual stimulation, and presentation of familiar and novel visual stimuli 24 hours after HFS. Power normalized as in A. Arrowhead indicates stimulus onset, white box indicates time window for assessment of change in oscillatory power (100-200 ms after stimulus reversal). D) Average +/- SEM change in oscillatory power, binned by frequency band during presentation of initial (black), familiar (red), and novel (blue) visual stimuli. HFS induces a significant decrease in average θ power in response to novel (blue) and familiar (red) visual stimulus, in all layers(RANOVA _(df, 2, 10)_, layer 2/3: F = 8.63, p = 0.002, layer 4: F = 8.47, p = 0.002, layer 5: F = 7.936, p = 0.003). Grey highlight = Bonferroni *post hoc* p < 0.05; n = 11 subjects.

### Visually-evoked reset of ongoing gamma oscillations predicts visual response potentiation

The phase of on-going LFP oscillations, and their reset in response to visual stimulation, can modify the amplitude of visually-evoked responses. To ask if visual response plasticity is coincident with visually-induced phase reset of the LFP we calculated the average ITPC during maximum visually-evoked activity (100-200 ms after stimulus onset). 24 hours after LFS, the familiar stimulus induced a significant increase in beta and gamma ITPC specifically in layer 4 (n = 16 subjects. RANOVA _(df, 2, 15)_, Bonferroni *post-hoc*, β: F = 5.20, p = 0.011; initial v. familiar: p = 0.045, γ: F = 8.82, p < 0.001; initial v. familiar: p = 0.005, Fig. 5A&B). In contrast, 24 hours after HFS, visual stimuli with familiar and novel orientations induced a significant increase in gamma ITPC in all cortical layers (n = 11 subjects. RANOVA _(df, 2, 10)_, Bonferroni *post-hoc*, layer 2/3: F = 19.42, p < 0.001; initial v. familiar: p = 0.001; initial v. novel: p = 0.003. Layer 4: F = 13.04, p < 0.001; initial v. familiar: p = 0.010; initial v. novel: p = 0.006. Layer 5: F = 5.29, p = 0.025; initial v. familiar: p = 0.022; initial v. novel: p = 0.021, Fig. 5C&D). Thus, the expression of increased gamma ITCP mirrored the locus and specificity of VEP potentiation in response to LFS and HFS.

**Figure 5.**
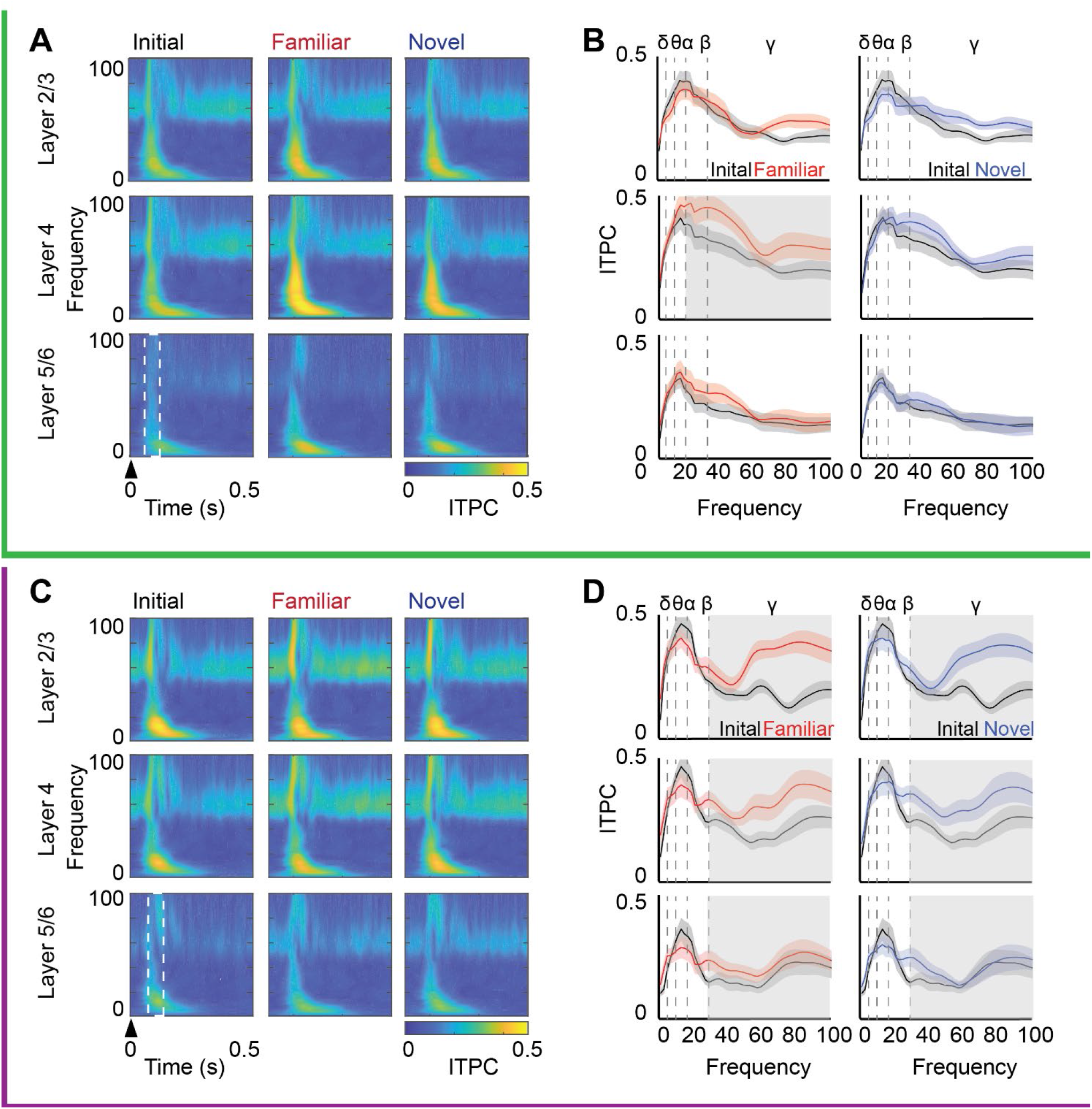
ITPC co-varies with visual response potentiation. Top, green, after LFS: A) Average inter-trial phase consistency 24 hours after LFS in response to initial, familiar and novel visual stimuli (ITPC; heat map) from 0-100 Hz (in 3 Hz bins; y axis) over time (x axis). Trial averaged from complex Morlet wavelet convolution. Arrowhead indicates stimulus onset; white box indicates time window for assessment of change in ITPC (100-200 ms after stimulus onset). B) Average ITPC power binned by frequency band (δ: 1-4, θ: 4-8, α: 8-13, β: 13-30, γ: 30-100Hz) in response to initial (black), familiar (red), and novel (blue) visual stimuli. 24 hours after LFS, the familiar visual stimulus induced a significant increase in layer 4 phase reset of β and γ oscillations relative to initial (black, RANOVA _(df, 2, 15)_, β: F = 5.20, p = 0.011, γ: F = 8.82, p < 0.001). Grey highlight = Bonferroni *post hoc* p < 0.05; n = 16 subjects. Bottom, purple, after HFS: C) Average ITPC 24 hours after HFS in response to initial, familiar and novel visual stimuli (ITPC; heat map) from 0-100 Hz (in 3 Hz bins; y axis) over time (x axis). Trial averaged from complex Morlet wavelet convolution. Arrowhead indicates stimulus onset; white box indicates time window for assessment of change in ITPC (100-200 ms after stimulus reversal). D) Average ITPC power binned by frequency band (δ: 1-4, θ: 4-8, α: 8-13, β: 13-30, γ: 30-100Hz) in response to initial (black), familiar (red), and novel (blue) visual stimuli. 24 hours after HFS, familiar (red) and novel (blue) visual stimuli induced a significant increase in γ oscillatory phase reset in all cortical layers relative to initial (black, RANOVA _(df, 2, 10)_, layer 2/3: F = 19.42, p < 0.001, layer 4: F = 13.04, p < 0.001, layer 5: F = 5.29, p = 0.025). Grey highlight = Bonferroni *post hoc* p < 0.05; n = 11 subjects.

To ask how changes in FS IN activity reflect the plasticity of visual responses, we examined the spike rate and oscillatory power of FS INs to visual stimuli with familiar (60°) and novel (150°) orientations 24 hours after LFS or HFS. Visual stimulation subsequent to HFS revealed a significant suppression of FS IN firing rates (n=11 subjects, 19 (initial), 20 (familiar), 20 (novel) units. One-way ANOVA _(df, 2, 57)_, Tukey *post-hoc*, F = 4.50, p = 0.015; initial v. familiar: p = 0.022; initial v. novel: p = 0.046. Fig. 6E) and a significant decrease the power of FS output at frequencies above theta (7 - 100Hz, n = 11 subjects, 19 (initial), 20 (familiar), 20 (novel) units. One-way ANOVA_(df, 2, 56)_, Bonferroni *post-hoc*, α: F = 6.862, p = 0.002, initial v. familiar, p = 0.004, initial v. novel, p = 0.011, β: F = 8.898, p < 0.001, initial v. familiar, p = 0.003, initial v. novel, p = 0.001, γ: F = 5.998, p = 0.004, initial v. familiar, p = 0.009, initial v. novel, p = 0.015; Fig. 6F). Suppression of FS IN firing rates and oscillatory power 24 hours after HFS was also observed in spontaneous activity prior to probe with familiar and novel stimuli (Fig. S2F). There were no significant changes in FS IN firing rate or power following LFS (n = 16 subjects, 22 (initial), 23 (familiar), 20 (novel) units; Fig. 6A&B).

**Figure 6.**
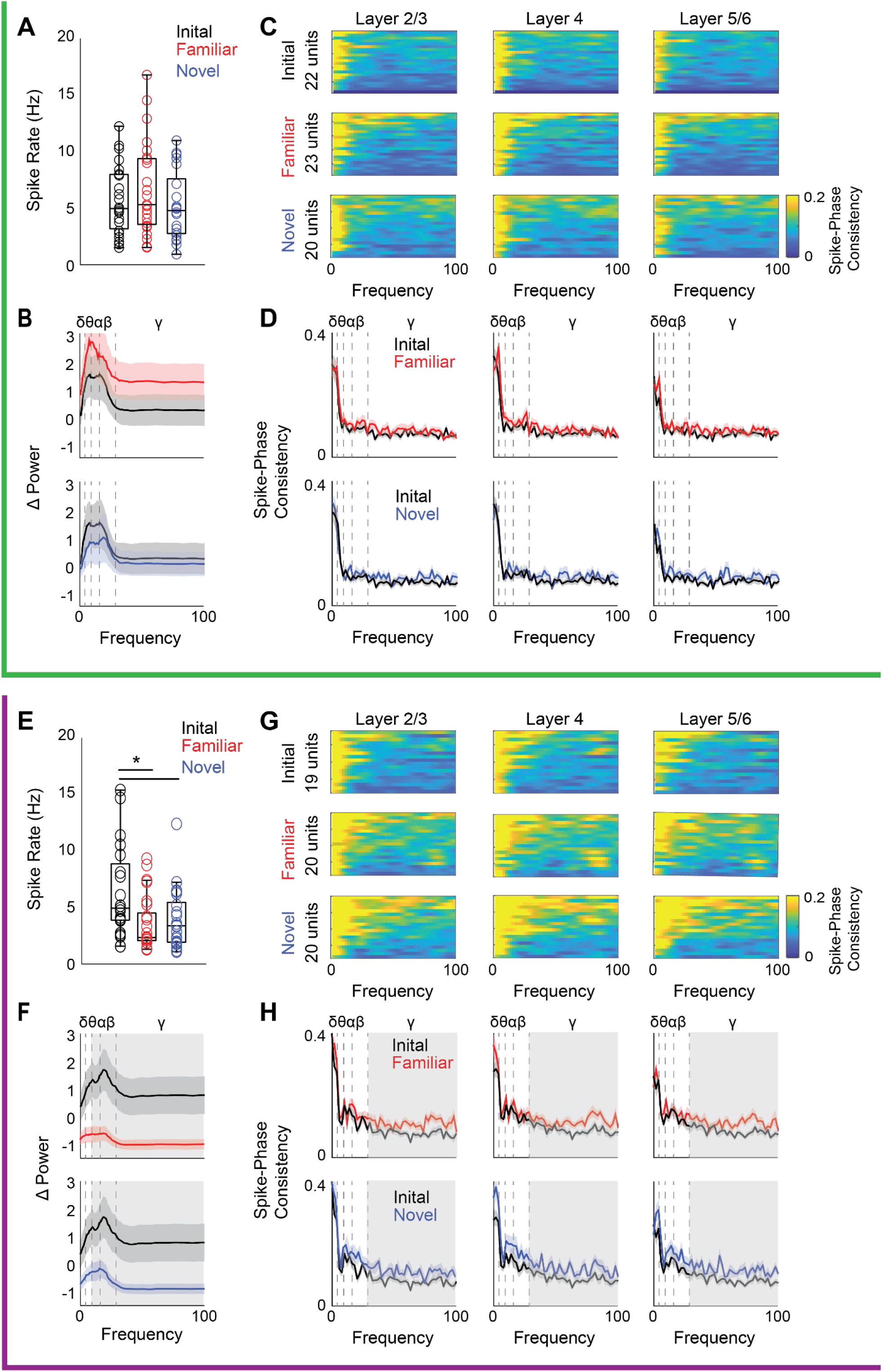
HFS decreases FS IN firing rates and power and increases FS IN spike-and LFP gamma phase consistency. Top, green, after LFS: A) No change in average spike rates of FS IN during presentation of familiar or novel visual stimuli 24 hours after LFS. B) No change in FS IN oscillatory power during presentation of familiar (red, top) or novel (blue, bottom) visual stimuli relative to initial (black). C) Heat map of spike-phase consistency of FS INs in response to initial, familiar and novel visual stimuli by cortical layer from 0 - 100 Hz during the 1000-200 ms following stimulus onset in subjects that received LFS. D) Average FS IN-LFP spike-phase consistency by cortical layer in response to initial (black), familiar (red), and novel (blue) stimuli. No significant difference in spike-phase consistency following LFS. Bottom, purple, after HFS: E) 24 hours after HFS, visual stimuli with familiar and novel orientations significantly decreased average FS IN spike rate (ANOVA _(df, 2, 56)_, F = 4.50, p = 0.015). * = Tukey *post hoc* p < 0.05; n = 11 subjects. F) Average change in FS IN power by frequency induced by initial (black) and familiar (red) visual stimuli. 24 hours after HFS, familiar and novel stimuli induced a significant decrease in the oscillatory power of FS IN across multiple frequencies (7-100Hz) compared to initial (black; ANOVA_(df, 2, 56)_, α: F = 6.862, p = 0.002, β: F = 8.898, p = 0.0004, γ: F = 5.998, p = 0.004). Grey highlight = Bonferroni *post hoc* p < 0.05, n = 11 subjects. G) Heat map of spike-phase consistency of FS IN during presentation of initial, familiar and novel stimuli by cortical layer from 0-100 Hz during the 100ms following stimulus reversal in subjects that received HFS. H) Average FS IN spike-LFP phase consistency by cortical layer induced by initial (black), familiar (red), and novel (blue) stimuli. 24 hours after HFS, familiar and novel visual stimuli induced a significant increase in FS IN spike-LFP γ phase consistency in all cortical layers (ANOVA_(df, 2, 56)_, Layer 2/3: F = 6.795, p = 0.002, layer 4: F = 5.655, p = 0.006, layer 5: F = 6.072, p = 0.004). Grey highlight = Bonferroni *post hoc* p < 0.05; n = 11 subjects.

We calculated spike-phase consistency between each FS IN and the LFP of each cortical layer to ask how FS IN firing is related to the phase of on-going LFP oscillatory activity. 24 hours after LFS there was no significant change in FS IN spike-LFP-phase consistency in any cortical layer in response to novel or familiar visual stimuli (n = 16 subjects, 22 (initial), 23 (familiar), 20 (novel) units; Fig. 6C&D). In contrast, 24 hours after HFS, presentation of visual stimuli with familiar and novel orientations induced a significant increase in FS IN spike-LFP phase coupling with gamma oscillations across all cortical layers (n = 11 subjects, 19 (initial), 20 (familiar), 20 (novel) units. One-way ANOVA_(df, 2, 56)_, Bonferroni *post-hoc*. Layer 2/3: F = 6.795, p = 0.002, initial v. familiar p = 0.009, initial v. novel p = 0.005. Layer 4: F = 5.655, p = 0.033, initial v. familiar p = 0.008, initial v. novel p = 0.005. Layer 5: F = 6.072, p = 0.004, initial v. familiar p = 0.026, initial v. novel p = 0.006; Fig. 6G&H). Thus, HFS induced a decrease in FS IN firing rates, and an increase in FS IN phase coupling with gamma, throughout the primary visual cortex.

### HFS enhances visual acuity

The increase in VEP magnitudes and ITPC following HFS was observed in response to visual stimuli with novel orientations and spatial frequencies, suggesting that HFS may induce a general enhancement of visual acuity. To test this prediction, we examined the impact of LFS and HFS on spatial acuity assessed by performance in a 2 alternative forced-choice spatial frequency detection task. Naïve mice (n = 12) were trained to associate a liquid reward with a simple visual stimulus (high contrast (100%), low spatial frequency (0.05 cpd, 45° sinusoidal grating; Fig. 7A). Subjects performed 30 trials per day, requiring 12.3 ± 1.05 days to reach the criterion of 25/30 correct trials (83%; Fig. 7B). To assess spatial acuity, the positive stimulus was rotated to a novel orientation (45°±15°), and subjects completed blocks of 10 trials with spatial frequencies from 0.05 cpd to 0.7 cpd in increments of 0.05 cpd. Spatial acuity was defined as highest spatial frequency with performance of ≥ 70% correct choices.

**Figure 7.**
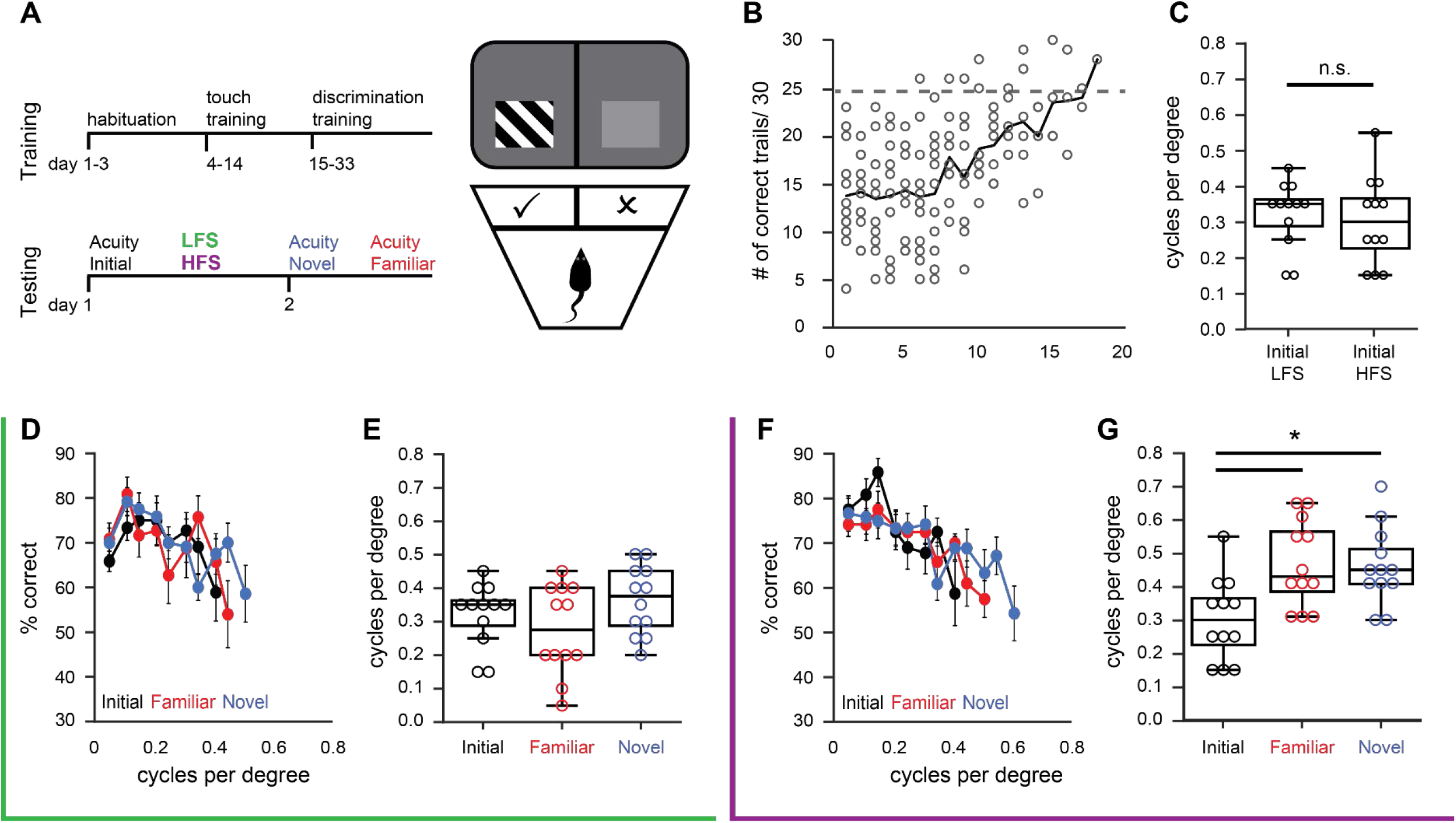
HFS enhances visual acuity. A) Left: timeline, subjects were trained in a 2-alternative forced choice visual detection task. Following training, baseline visual acuity was assessed using a novel stimulus orientation and followed by LFS or HFS. Spatial acuity was tested again 24 hour later. Right: cartoon depicting modified Bussey chamber with plexiglass divider introducing a choice point for the assessment of stimulus spatial frequency. B) All subjects reached task criterion of 25/30 correct trials by 18 days of training. C) No significant difference in initial visual acuity prior to the delivery of LFS or HFS (Student’s t-test, p = 0.63). D) Average frequency of seeing curves for initial (black), novel (blue) and familiar (red) visual stimulus orientations in subjects that received LFS (green box). E) No significant difference in spatial acuity probed with initial (black, before LFS), novel (blue, after LFS) and familiar (red, after LFS) visual stimulus orientations (n = 12). F) Average frequency of seeing curves for initial (black), novel (blue) and familiar (red) visual stimulus orientations in subjects that received HFS (purple box). G) Spatial acuity probed with novel (blue, after HFS) and familiar (red, after HFS) visual stimulus orientations was significantly increased following HFS (black, before HFS; RANOVA _(df, 2, 11)_, F = 6.817, p = 0.005). * = Bonferroni *post hoc* p < 0.05; n = 12.

Following determination of baseline spatial acuity, subjects passively viewed 200 x 1 second trials of either LFS or HFS at a novel orientation (45° ± 30°). There was no significant difference in baseline spatial acuity between subjects assigned to LFS and HFS groups (average +/- SEM cpd, LFS; 0.33 ± 0.02; HFS 0.30 ± 0.03 cpd, Student’s t-test, p = 0.63,Fig. 6C). 24 hours following visual stimulation, visual performance was assessed at the familiar (45° ± 30°) or another novel orientation (45° ± 60°). Following LFS, we observed no change in spatial acuity assessed with the familiar (0.30 ± 0.03 cpd) or novel visual stimulus (0.37 ± 0.03 cpd; n = 12, RANOVA _(df, 2, 11)_, F = 4.02, p = 0.032, initial v familiar p = 0.303, initial v. novel p = 0.303; Fig. 7D&E). In contrast, 24 hours after HFS, spatial acuity was significantly enhanced in response to familiar (0.46 ± 0.03 cpd) and novel visual stimulus orientations (0.46 ± 0.03; n = 12, RANOVA _(df, 2, 11)_, Bonferroni *post-hoc*, F = 6.87, p = 0.005; initial v. familiar, p = 0.011; initial v. novel, p = 0.011; Fig. 7F&G). Together this demonstrates that HFS induced a sustained, and highly generalizable increase in visual function.

## Discussion

We used chronic electrophysiological recordings in the visual cortex of awake mice and a psychophysical measurement of visual acuity to delineate the distinct cellular, circuit, and perceptual responses to high (HFS) and low (LFS) frequency repetitive visual stimulation. We demonstrate that repetitive high and low frequency visual stimulation induced experience-dependent plasticity of visual responses with distinct loci of expression. A single, short bout of LFS (1s x 200 presentations) was sufficient to engage response potentiation of layer 4 VEPs, with similar selectivity for the familiar visual stimulus but smaller magnitude than response potentiation induced by daily LFS (Frenkel et al., 2006). In contrast, a single bout of HFS was sufficient to induce potentiation of VEP amplitudes in all layers of V1, that generalizes to novel stimuli. Single unit recordings reveal that HFS selectively suppressed the output of FS INs V1. Furthermore, HFS primed high frequency (gamma) cortical oscillations of the LFP for visually-evoked phase rest and synchronized fast-spiking interneuron output with gamma rhythms. These long-lasting changes in cortical dynamics, maintained 24 hours after HFS, is revealed as response potentiation to all subsequent visual stimuli and enhanced visual acuity.. Together this demonstrates that the temporal parameter of repetitive visual stimulation is a decisive factor in the regulation of visual response plasticity.

It is increasingly appreciated that neurons in V1 reflect more than visual input. Indeed neuronal activity patterns are altered by locomotion (Niell and Stryker, 2010), generalized motor movements, arousal state (Stringer et al., 2019) and input from other sensory systems (Iurilli et al., 2012). Similarly, ongoing cortical rhythms in V1, including beta (12 - 30 Hz) and gamma (30 - 100 Hz), are regulated by visual inputs including contrast intensity (Saleem et al., 2017), visual stimulus size (Veit et al., 2017), and non-visual inputs, including the encoding and memory of complex temporal sequences (Gavornik and Bear 2014; Han et al., 2008). For example, reward timing is encoded by an increase in the power of 5 - 10 Hz (theta) frequency oscillations in V1 (Zold and Hussain Shuler, 2015). Similarly, repetitive visual stimulation with a familiar stimulus generates an increase in the visually-evoked power of theta (4 - 8 Hz), alpha (8 - 12 Hz) and beta (12 - 30 Hz) oscillations in layer 4 of V1 (Kissinger et al., 2018; Kissinger et al., 2020). Following a single bout of LFS, we observed a similar increase in mid-range oscillatory activity in the visual cortex in response to a familiar visual stimulus. However, following HFS, theta power was decreased across all cortical layers in response to familiar and novel visual stimulus orientations and spatial frequencies. The latter is consistent with the observation that HFS suppresses the firing rate of FS INs, as FS INs have been shown to drive the production of theta oscillations within the cortex and hippocampus (Buzsáki, 2002; Stark et al., 2013). In addition we saw a decrease in low frequency power (1 – 8 Hz) in all cortical layers in response to stimuli with novel and familiar orientations, as seen during directed attention (Schroeder and Lakatos, 2009).

However, changes in power of cortical oscillations in the absence of synchronization of oscillatory phase could be expected to reduce response magnitude by increasing variability, while phase synchronization with incoming stimulation would increase response magnitude and decrease response variability. Indeed, we found that visually-induced phase reset of cortical gamma oscillations predicts both the location and specificity of visual response potentiation in response to LFS and HFS. Familiar and novel visual stimuli subsequent to HFS reset the phase of ongoing gamma oscillations throughout V1, while visually-induced gamma phase reset following LFS was expressed in layer 4 in response to the familiar visual stimulus. The sensitivity of gamma oscillations for visually-evoked phase reset following HFS likely reflects the specifics of generation and modulation of high frequency oscillations. Cortical gamma oscillations are generated by feedforward thalamo-cortical connections, while phase synchrony is modulated/ entrained by the output of cortical parvalbumin expressing FS INs (Bastos et al., 2015; van Kerkoerle et al., 2014; Saleem et al., 2017; Spaak et al., 2012; Cardin et al., 2009; Chen et al., 2017; Sohal et al., 2009). A reduced influence of FS IN output on gamma following HFS could promote enhanced sensitivity to gamma phase reset by visual stimulation and enhanced generalizability of response potentiation. The generalizability of response potentiation following HFS mirrors previous reports that optogenetic suppression of FS IN output decreases the selectivity of visual response potentiation induced by daily LFS (Kaplan et al., 2016).

LFS and HFS will recruit activity in different subsets of neurons in mouse V1, tuned to lower and higher temporal frequencies respectively (Gao et al., 2010). Nonetheless, the temporal frequency of the initial visual stimulation was reflected in changes in LFP oscillatory power and phase reset, and FS IN output. The temporal frequency of our high frequency stimulation (20 Hz) is close to flicker fusion in the murine visual system, and may therefore drive the largest number of pyramidal neurons to spike at high frequency (Durand et al., 2016; Tanimoto et al., 2015). Suppression of FS IN firing rates or changes in synchrony of FS IN spiking with oscillations were not observed during the HFS, suggesting additional processes are necessary for the expression observed at 24 hours. One likely possibility is that sleep is necessary to consolidate long term changes in cortical activity induced by HFS, as has been shown for the selective response potentiation following daily LFS (Aton et al., 2014). HFS induced generalized plasticity of visual responses throughout V1 and a parallel increase in spatial acuity revealed by visual task performance. However, we cannot rule out the possibility that the synaptic changes underlying improved spatial acuity following HFS reflect plasticity beyond V1, such as the latero-intermediate area, where neurons are tuned to higher temporal and spatial frequencies than V1 (Marshel et al., 2011). Interestingly, the perceptual habituation induced by daily repetition of low frequency stimulation transfers from training to testing environments and shares stimulus selectivity and NMDAR-dependence with coincident visual response potentiation, indicating a shared synaptic locus (Cooke et al., 2015).

Our findings provide mechanistic insight into the cellular, circuit and perceptual response of mouse V1 to high (HFS) and low (LFS) frequency repetitive visual stimulation, which may lend insight into sensory system plasticity in other species. In humans, the temporal frequency-dependence of visual perception plasticity is well documented. A high frequency (20 Hz) luminance flicker induces a long-lasting improvement in luminance change detection, while a low frequency (1 Hz) luminance flicker briefly reduces performance (Beste et al., 2011). Similarly, high frequency (20Hz), but not low frequency (1 Hz), presentation of an oriented bar or a flickering sinusoidal grating improves orientation discrimination (Marzoll et al., 2018). High temporal frequency visual stimulation (10 Hz) during a word recognition task improved word recall, demonstrating that repetitive visual stimulation can impact the function of neural circuitry outside of primary visual cortex (Williams, 2001; Williams et al., 2006). Performance on sensory and memory tasks are also manipulated by trans-cranial brain stimulation techniques. Indeed, low frequency rTMS (1-3 Hz pulses) stimulation over the visual cortex in cats induced transient depression of the amplitude of visual response amplitudes, while high frequency stimulation (10 Hz) transiently potentiated visual responses (Aydin-Abidin et al., 2006). The HFS visual stimulation protocol described here results in long-lasting and highly generalizable enhancement of visual function and may therefore be an attractive candidate for non-invasive vision therapy.

## Acknowledgements

This work was supported by the National Institutes of Health grant EY016431 and EY025922 to EMQ.

---

**Figure S1.**
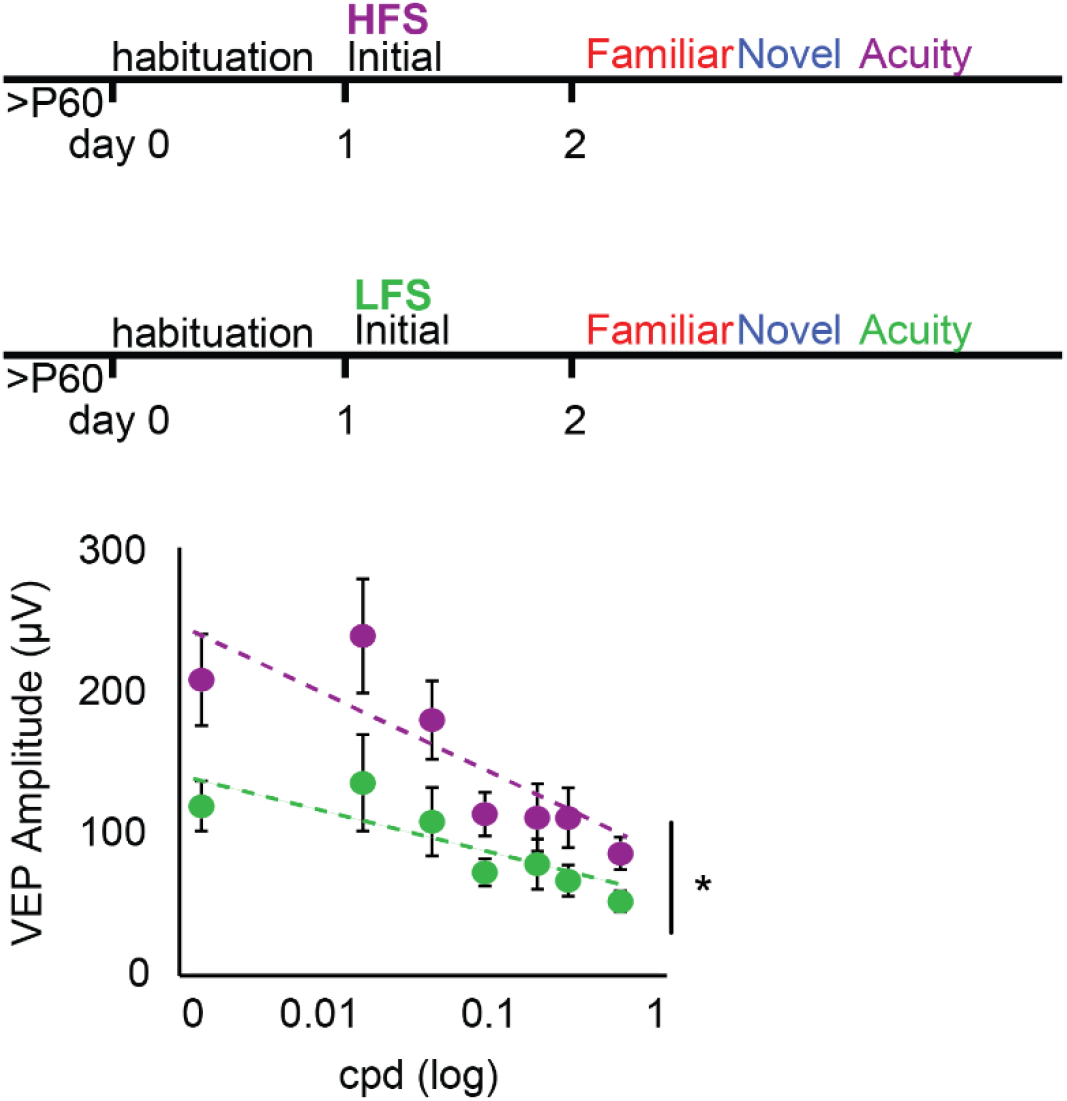
HFS enhances VEP amplitudes evoked by novel spatial frequencies. Following HFS, a significant increase in layer 4 VEP amplitudes is observed in response to visual stimuli with a novel orientation, across a range of spatial frequencies (purple) compared to LFS. (Between groups RANOVA_(df,6,1)_, F = 5.88, p = 0.035). * = p < 0.05; n = 9 (HFS), 6 (LFS) subjects.

**Figure S2.**
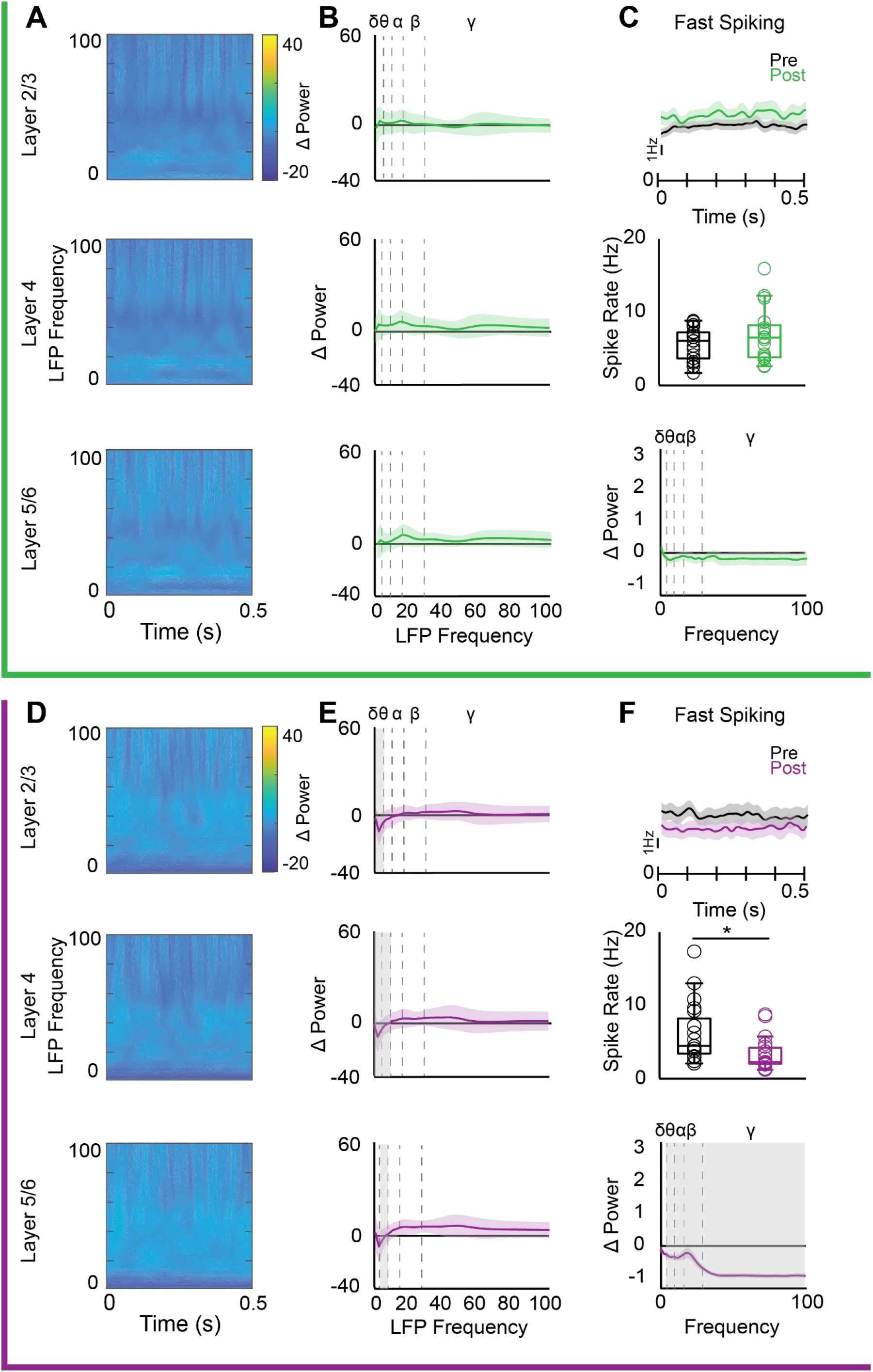
Distinct sustained effects of LFS and HFS on spontaneous oscillatory power and FS IN spiking. Top, green, after LFS: A) Heat map, average change in spontaneous oscillatory power 24 hours after LFS (% change from pre-stimulation spontaneous activity) from 0-100 Hz (3 Hz bins; y axis) over time (x axis) by cortical layer. B) Average % change in spontaneous oscillatory power 24 hours after LFS, binned by frequency band (δ: 1-4, θ: 4-8, α: 8-13, β: 13-30, γ: 30-100 Hz). No change in spontaneous oscillatory power 24 hours after LFS in any cortical layer (n = 16). C) Top, average +/- SEM time-histogram of FS INs, pre (black) and 24 hours post LFS (green). Middle, average FS IN spike rates pre (black) and 24 hours post LFS (green). Bottom, no change in oscillatory power of FS IN spontaneous activity 24 hours post LFS, binned by frequency band. Significant decrease in θ, β, and γ oscillatory power 24 hours after LFS. Bottom, purple, after HFS: D) Heat map, average change in spontaneous oscillatory power 24 hours after HFS (% change from pre-HFS spontaneous activity) from 0 to 100 Hz (3 Hz bins; y axis) over time (x axis) by cortical layer. E) Average % change in spontaneous oscillatory power 24 hours after HFS, binned by frequency band. 24 hours after HFS, δ power is significantly decreased in all layers (one sided t-test, grey highlight = p < 0.05; n = 11). F) Top, average +/- SEM time-histogram of FS INs pre LFS (blank) and 24 hours post HFS (purple). Middle, Significant decrease in average FS IN firing rates 24 hours after HFS (unpaired t-test, * = p < 0.05; n = 19 (pre), 18 (post) units, 11 subjects). Bottom, average change in spontaneous oscillatory power of FS INs 24 hours post LFS, binned by frequency band. Significant decrease in all frequency bands above δ 24 hours after HFS (One-sided t-test, grey highlight = p < 0.05; n = 18 units, 11 subjects).

## References

Aton, S.J., Suresh, A., Broussard, C., and Frank, M.G. (2014). Sleep Promotes Cortical Response Potentiation Following Visual Experience. Sleep 37, 1163–1170.

Aydin-Abidin, S., Moliadze, V., Eysel, U.T., and Funke, K. (2006). Effects of repetitive TMS on visually evoked potentials and EEG in the anaesthetized cat: dependence on stimulus frequency and train duration. J. Physiol. 574, 443–455.

Bastos, A.M., Vezoli, J., Bosman, C.A., Schoffelen, J.-M., Oostenveld, R., Dowdall, J.R., De Weerd, P., Kennedy, H., and Fries, P. (2015). Visual Areas Exert Feedforward and Feedback Influences through Distinct Frequency Channels. Neuron 85, 390–401.

Beste, C., Wascher, E., Güntürkün, O., and Dinse, H.R. (2011). Improvement and Impairment of Visually Guided Behavior through LTP- and LTD-like Exposure-Based Visual Learning. Curr. Biol. 21, 876–882.

Bliss, T.V.P., and Lømo, T. (1973). Long-lasting potentiation of synaptic transmission in the dentate area of the anaesthetized rabbit following stimulation of the perforant path. J. Physiol. 232, 331–356.

Brainard, D.H. (1997). The Psychophysics Toolbox. Spat. Vis. 10, 433–436.

Brickwedde, M., Krüger, M.C., and Dinse, H.R. (2019). Somatosensory alpha oscillations gate perceptual learning efficiency. Nat. Commun. 10, 263.

Brickwedde, M., Schmidt, M.D., Krüger, M.C., and Dinse, H.R. (2020). 20 Hz Steady-State Response in Somatosensory Cortex During Induction of Tactile Perceptual Learning Through LTP-Like Sensory Stimulation. Front. Hum. Neurosci. 14, 257.

Bridi, M.C.D., de Pasquale, R., Lantz, C.L., Gu, Y., Borrell, A., Choi, S.-Y., He, K., Tran, T., Hong, S.Z., Dykman, A., et al. (2018). Two distinct mechanisms for experience-dependent homeostasis. Nat. Neurosci. 21, 1.

Buzsáki, G. (2002). Theta oscillations in the hippocampus. Neuron 33, 325–340.

Cardin, J.A., Carlén, M., Meletis, K., Knoblich, U., Zhang, F., Deisseroth, K., Tsai, L.-H., and Moore, C.I. (2009). Driving fast-spiking cells induces gamma rhythm and controls sensory responses. Nature 459, 663–667.

Chang, M.C., Park, J.M., Pelkey, K.A., Grabenstatter, H.L., Xu, D., Linden, D.J., Sutula, T.P., McBain, C.J., and Worley, P.F. (2010). Narp regulates homeostatic scaling of excitatory synapses on parvalbumin-expressing interneurons. Nat. Neurosci. 13, 1090–1097.

Chen, G., Zhang, Y., Li, X., Zhao, X., Ye, Q., Lin, Y., Tao, H.W., Rasch, M.J., and Zhang, X. (2017). Distinct Inhibitory Circuits Orchestrate Cortical beta and gamma Band Oscillations. Neuron 96, 1403–1418.e6.

Clapp, W.C., Eckert, M.J., Teyler, T.J., and Abraham, W.C. (2006). Rapid visual stimulation induces N-methyl-D-aspartate receptor-dependent sensory long-term potentiation in the rat cortex. Neuroreport 17, 511–515.

Cooke, S.F., and Bear, M.F. (2010). Visual experience induces long-term potentiation in the primary visual cortex. J. Neurosci. 30, 16304–16313.

Cooke, S.F., Komorowski, R.W., Kaplan, E.S., Gavornik, J.P., and Bear, M.F. (2015). Visual recognition memory, manifested as long-term habituation, requires synaptic plasticity in V1. Nat. Neurosci. 18, 262–271.

Dinse, H.R., Ragert, P., Pleger, B., Schwenkreis, P., and Tegenthoff, M. (2003). Pharmacological modulation of perceptual learning and associated cortical reorganization. Science (80-.). 301, 91–94.

Dudek, S.M., and Bear, M.F. (1992). Homosynaptic long-term depression in area CA1 of hippocampus and effects of N-methyl-D-aspartate receptor blockade. Proc. Natl. Acad. Sci. U. S. A. 89, 4363–4367.

Durand, S., Iyer, R., Mizuseki, K., De Vries, S., Mihalas, S., and Reid, R.C. (2016). A comparison of visual response properties in the lateral geniculate nucleus and primary visual cortex of awake and anesthetized mice. J. Neurosci. 36, 12144–12156.

Fiebelkorn, I.C., Pinsk, M.A., and Kastner, S. (2018). A Dynamic Interplay within the Frontoparietal Network Underlies Rhythmic Spatial Attention. Neuron 99, 842–853.e8.

Frenkel, M.Y., Sawtell, N.B., Diogo, A.C., Yoon, B., Neve, R.L., and Bear, M.F. (2006). Instructive effect of visual experience in mouse visual cortex. Neuron 51, 339–349.

Gao, E., DeAngelis, G.C., and Burkhalter, A. (2010). Parallel input channels to mouse primary visual cortex. J. Neurosci. 30, 5912–5926.

Gavornik, J.P., and Bear, M.F. (2014). Learned spatiotemporal sequence recognition and prediction in primary visual cortex. Nat. Neurosci. 17, 732–737.

Gu, Y., Huang, S., Chang, M.C., Worley, P., Kirkwood, A., and Quinlan, E.M. (2013). Obligatory role for the immediate early gene NARP in critical period plasticity. Neuron 79, 335.

Guo, W., Clause, A.R., Barth-Maron, A., and Polley, D.B. (2017). A Corticothalamic Circuit for Dynamic Switching between Feature Detection and Discrimination. Neuron 95, 180–194.e5.

Han, F., Caporale, N., and Dan, Y. (2008). Reverberation of Recent Visual Experience in Spontaneous Cortical Waves. Neuron 60, 321–327.

Horner, A.E., Heath, C.J., Hvoslef-Eide, M., Kent, B.A., Kim, C.H., Nilsson, S.R.O., Alsiö, J., Oomen, C.A., Holmes, A., Saksida, L.M., et al. (2013). The touchscreen operant platform for testing learning and memory in rats and mice. Nat. Protoc. 8, 1961–1984.

Howe, W.M., Gritton, H.J., Lusk, N.A., Roberts, E.A., Hetrick, V.L., Berke, J.D., and Sarter, M. (2017). Acetylcholine Release in Prefrontal Cortex Promotes Gamma Oscillations and Theta-Gamma Coupling during Cue Detection. J. Neurosci. 37, 3215–3230.

Iurilli, G., Ghezzi, D., Olcese, U., Lassi, G., Nazzaro, C., Tonini, R., Tucci, V., Benfenati, F., and Medini, P. (2012). Sound-Driven Synaptic Inhibition in Primary Visual Cortex. Neuron 73, 814–828.

Jutras, M.J., Fries, P., and Buffalo, E.A. (2009). Gamma-band synchronization in the macaque hippocampus and memory formation. J. Neurosci. 29, 12521–12531.

Kaplan, E.S., Cooke, S.F., Komorowski, R.W., Chubykin, A.A., Thomazeau, A., Khibnik, L.A., Gavornik, J.P., and Bear, M.F. (2016). Contrasting roles for parvalbumin-expressing inhibitory neurons in two forms of adult visual cortical plasticity. Elife 5.

van Kerkoerle, T., Self, M.W., Dagnino, B., Gariel-Mathis, M.-A., Poort, J., van der Togt, C., and Roelfsema, P.R. (2014). Alpha and gamma oscillations characterize feedback and feedforward processing in monkey visual cortex. Proc. Natl. Acad. Sci. U. S. A. 111, 14332–14341.

Kim, Y.J., Grabowecky, M., Paller, K.A., Muthu, K., and Suzuki, S. (2007). Attention induces synchronization-based response gain in steady-state visual evoked potentials. Nat. Neurosci. 10, 117–125.

Kirkwood, A., Rioult, M.G., and Bear, M.F. (1996). Experience-dependent modification of synaptic plasticity in visual cortex. Nature 381, 526–528.

Kissinger, S.T., Pak, A., Tang, Y., Masmanidis, S.C., and Chubykin, A.A. (2018). Oscillatory Encoding of Visual Stimulus Familiarity. J. Neurosci. 38, 6223–6240.

Kissinger, S.T., Wu, Q., Quinn, C.J., Anderson, A.K., Pak, A., and Chubykin, A.A. (2020). Visual Experience-Dependent Oscillations and Underlying Circuit Connectivity Changes Are Impaired in Fmr1 KO Mice. Cell Rep. 31, 107486.

Larson, J., Wong, D., and Lynch, G. (1986). Patterned stimulation at the theta frequency is optimal for the induction of hippocampal long-term potentiation. Brain Res. 368, 347–350.

Lisman, J. (2010). Working Memory: The Importance of Theta and Gamma Oscillations. Curr. Biol. 20, R490–R492.

Mayford, M., Wang, J., Kandel, E.R., and O’Dell, T.J. (1995). CaMKII regulates the frequency-response function of hippocampal synapses for the production of both LTD and LTP. Cell 81, 891–904.

Marshel, J.H., Garrett, M.E., Nauhaus, I., and Callaway, E.M. (2011). Functional specialization of seven mouse visual cortical areas. Neuron 72, 1040–1054.

Marzoll, A., Saygi, T., and Dinse, H.R. (2018). The effect of LTP- and LTD-like visual stimulation on modulation of human orientation discrimination. Sci. Rep. 8, 16156.

Mitzdorf, U. (1985). Current source-density method and application in cat cerebral cortex: investigation of evoked potentials and EEG phenomena. Physiol. Rev. 65, 37–100.

Montgomery, S.M., and Buzsáki, G. (2007). Gamma oscillations dynamically couple hippocampal CA3 and CA1 regions during memory task performance. Proc. Natl. Acad. Sci. U. S. A. 104, 14495–14500.

Murase, S., Lantz, C.L., and Quinlan, E.M. (2017). Light reintroduction after dark exposure reactivates plasticity in adults via perisynaptic activation of MMP-9. Elife 6.

Nabavi, S., Fox, R., Proulx, C.D., Lin, J.Y., Tsien, R.Y., and Malinow, R. (2014). Engineering a memory with LTD and LTP. Nature 511, 348–352.

Niell, C.M., and Stryker, M.P. (2008). Highly selective receptive fields in mouse visual cortex. J. Neurosci. 28, 7520–7536.

Niell, C.M., and Stryker, M.P. (2010). Modulation of visual responses by behavioral state in mouse visual cortex. Neuron 65, 472–479.

O’Brien, R.J., Xu, D., Petralia, R.S., Steward, O., Huganir, R.L., and Worley, P. (1999). Synaptic clustering of AMPA receptors by the extracellular immediate-early gene product Narp. Neuron 23, 309–323.

O’Riordan, K.J., Hu, N.W., and Rowan, M.J. (2018). Physiological activation of mGlu5 receptors supports the ion channel function of NMDA receptors in hippocampal LTD induction in vivo. Sci. Rep. 8, 1–12

ten Oever, S., Schroeder, C.E., Poeppel, D., Van Atteveldt, N., Mehta, A.D., Mégevand, P., Groppe, D.M., and Zion-Golumbic, E. (2017). Low-frequency cortical oscillations entrain to subthreshold rhythmic auditory stimuli. J. Neurosci. 37, 4903–4912.

Park, H., Lee, D.S., Kang, E., Kang, H., Hahm, J., Kim, J.S., Chung, C.K., Jiang, H., Gross, J., and Jensen, O. (2016). Formation of visual memories controlled by gamma power phase-locked to alpha oscillations. Sci. Rep. 6, 28092.

Pegado, F., Vankrunkelsven, H., Steyaert, J., Boets, B., De Beeck, H.O., and Op de Beeck, H. (2016). Exploring the Use of Sensorial LTP/LTD-Like Stimulation to Modulate Human Performance for Complex Visual Stimuli. PNAS 11, e0158312.

Pelli, D.G. (1997). The VideoToolbox software for visual psychophysics: transforming numbers into movies. Spat. Vis. 10, 437–442.

Poggio, T., Fahle, M., Edelman, S., Askenasy, J., and Sagi, D. (1992). Fast perceptual learning in visual hyperacuity. Science (80-.). 256, 1018–1021.

Rodríguez-Durán, L.F., Martínez-Moreno, A., and Escobar, M.L. (2017). Bidirectional modulation of taste aversion extinction by insular cortex LTP and LTD. Neurobiol. Learn. Mem. 142, 85–90.

Ross, R.M., McNair, N.A., Fairhall, S.L., Clapp, W.C., Hamm, J.P., Teyler, T.J., and Kirk, I.J. (2008). Induction of orientation-specific LTP-like changes in human visual evoked potentials by rapid sensory stimulation. Brain Res. Bull. 76, 97–101

Saleem, A.B., Lien, A.D., Krumin, M., Haider, B., Rosón, M.R., Ayaz, A., Reinhold, K., Busse, L., Carandini, M., Harris, K.D., et al. (2017). Subcortical Source and Modulation of the Narrowband Gamma Oscillation in Mouse Visual Cortex. Neuron 93, 315–322.

Sanders, P.J., Thompson, B., Corballis, P.M., Maslin, M., and Searchfield, G.D. (2018). A review of plasticity induced by auditory and visual tetanic stimulation in humans. Eur. J. Neurosci. 48, 2084–2097.

Sauseng, P., Klimesch, W., Gruber, W.R.R., Hanslmayr, S., Freunberger, R., and Doppelmayr, M. (2007). Are event-related potential components generated by phase resetting of brain oscillations? A critical discussion. Neuroscience 146, 1435–1444.

Schroeder, C.E., and Lakatos, P. (2009). Low-frequency neuronal oscillations as instruments of sensory selection. Trends Neurosci. 32, 9–18

Sohal, V.S., Zhang, F., Yizhar, O., and Deisseroth, K. (2009). Parvalbumin neurons and gamma rhythms enhance cortical circuit performance. Nature 459, 698–702.

Spaak, E., Bonnefond, M., Maier, A., Leopold, D.A., and Jensen, O. (2012). Layer-Specific Entrainment of Gamma-Band Neural Activity by the Alpha Rhythm in Monkey Visual Cortex. Curr. Biol. 22, 2313–2318.

Stark, E., Eichler, R., Roux, L., Fujisawa, S., Rotstein, H.G., and Buzsáki, G. (2013). Inhibition-Induced theta resonance in cortical circuits. Neuron 80, 1263–1276.

Stringer, C., Pachitariu, M., Steinmetz, N., Reddy, C.B., Carandini, M., and Harris, K.D. (2019). Spontaneous behaviors drive multidimensional, brainwide activity. Science 364, 255.

Tanimoto, N., Sothilingam, V., Kondo, M., Biel, M., Humphries, P., and Seeliger, M.W. (2015). Electroretinographic assessment of rod- and cone-mediated bipolar cell pathways using flicker stimuli in mice. Sci. Rep. 5, 10731.

Veit, J., Hakim, R., Jadi, M.P., Sejnowski, T.J., and Adesnik, H. (2017). Cortical gamma band synchronization through somatostatin interneurons. Nat. Neurosci. 20, 951–959.

Williams, J.. (2001). Frequency-specific effects of flicker on recognition memory. Neuroscience 104, 283–286.

Williams, J., Ramaswamy, D., and Oulhaj, A. (2006). 10 Hz flicker improves recognition memory in older people. BMC Neurosci. 7, 21.

Xing, D., Yeh, C.I., Burns, S., and Shapley, R.M. (2012). Laminar analysis of visually evoked activity in the primary visual cortex. Proc. Natl. Acad. Sci. U. S. A. 109, 13871–13876.

Xu, D., Hopf, C., Reddy, R., Cho, R.W., Guo, L., Lanahan, A., Petralia, R.S., Wenthold, R.J., O’Brien, R.J., and Worley, P. (2003). Narp and NP1 Form Heterocomplexes that Function in Developmental and Activity-Dependent Synaptic Plasticity. Neuron 39, 513–528.

Zaehle, T., Clapp, W.C., Hamm, J.P., Meyer, M., and Kirk, I.J. (2007). Induction of LTP-like changes in human auditory cortex by rapid auditory stimulation: An FMRI study. Restor. Neurol. Neurosci. 25, 251–259.

Zold, C.L., and Hussain Shuler, M.G. (2015). Theta Oscillations in Visual Cortex Emerge with Experience to Convey Expected Reward Time and Experienced Reward Rate. J. Neurosci. 35, 9603–9614.

